# Liver ultrasound elastography for the evaluation of periportal fibrosis in schistosomiasis mansoni: A cross-sectional study

**DOI:** 10.1101/450494

**Authors:** Joelma Carvalho Santos, Andrea Dória Batista, Carla Maria Mola Vasconcelos, Roberto Souza Lemos, Valter Romão de Souza, Alain Dessein, Hélia Dessein, Silvia Maria Lucena Montenegro, Edmundo Pessoa Almeida Lopes, Ana Lúcia Coutinho Domingues

## Abstract

**Background:** ARFI elastrography has been used as a noninvasive method to assess the severity of liver fibrosis in viral hepatitis, although with few studies in schistosomiasis mansoni. We aimed to evaluate the performance of point shear wave elastography (pSWE) for predicting significant periportal fibrosis (PPF) in schistosomotic patients and to determine its best cutoff point.

**Methodology/Principal findings:** This cross-sectional study included 358 adult schistosomotic patients subjected to US and pSWE on the right lobe. Two hundred two patients (62.0%) were women, with a median age of 54 (ranging 18-92) years. The pSWE measurements were compared to the US patterns of PPF, as gold standard, according to the Niamey classification. The performance of pSWE was calculated as the area under the ROC curve (AUC). Patients were further classified into two groups: 86 patients with mild PPF and 272 patients with significant PPF. The median pSWE of the significant fibrosis group was higher (1.40 m/s) than that of mild fibrosis group (1.14 m/s, p<0.001). AUC was 0.719 with ≤1.11 m/s as the best cutoff value for excluding significant PPF. Sensitivity and negative predictive values were 80.5% and 40.5%, respectively. Whereas, for confirming significant PPF, the best cutoff value was >1.39 m/s, with specificity of 86.1% and positive predictive value of 92.0%.

**Conclusions/Significance:** pSWE was able to differentiate significant from mild PPF, with better performance to predict significant PPF.

**Author summary:** In the developing world, over 207 million people are infected with parasitic Schistosoma worms. Among the species of Schistosoma that infect humans Schistosoma mansoni is one of the most common causes of illness. Here, we investigated the performance of point shear wave elastography (pSWE) for predicting significant periportal fibrosis (PPF) in schistosomotic patients and to determine its best cutoff point. We examined 358 people from northeast of Brazil for Schistosoma infections. The present study showed that pSWE was able to differentiate significant from mild PPF, with better performance to predict significant PPF.

## Introduction

Schistosomiasis is considered a public health problem in many parts of the world and one of the most frequent causes of liver fibrosis worldwide [1]. In Brazil, schistosomiasis mansoni infection affects 1.5 millions of people, particularly in the Northeastern region [2]. The state of Pernambuco is considered an endemic area with 4207 cases in 2016, and a mortality average of 173 deaths per year between 2005 and 2015 [3].

Hepatic schistosomiasis represents the best known form of chronic disease with a wide range of clinical manifestations [4]. Accordingly, periportal fibrosis (PPF) or Symmers fibrosis is a schistosoma-induced liver disease, and the progression of PPF from hepatointestinal (HIS) to hepatosplenic (HSS) form is associated with the development of portal hypertension and its complications, such as esophageal varices and hypertensive gastropathy. Moreover, the patients with significant PPF consequently presents more portal hypertension and more risk of upper gastrointestinal bleeding [5].

The gold standard method in evaluating schistosomal hepatic fibrosis was wedge liver biopsy, a safe procedure performed during abdominal surgery, but not justified in non-surgical patients. An alternative procedure, percutaneous liver biopsy, has low sensitivity because it retrieves small and fragmented samples with few portal tracts [6]. Furthermore, liver biopsy is invasive, usually unsuitable for long-term follow-up and not applicable in the field, indicating that there is no gold standard for schistosomal portal fibrosis available [7]. Therefore, imaging techniques, such as ultrasound (US) scan, have been proven valuable for liver morbidity assessment and its complications in endemic areas [1,8,9].

Hence, US is considered a complementary tool often used to assist the diagnosis of the *S. mansoni* infection, mainly on the study of the liver damage caused by the chronic infection [10]. US also assess liver, spleen and portal vein size, and it is important to exclude or confirm other intra-abdominal diseases [11]. For the assessment of PPF, and consequently schistosomiasis-related morbidity, US is one of the most important means for this purpose, especially with the formulation of a standardized WHO Niamey-Belo Horizonte protocol. [1,12]. The WHO Niamey-Belo Horizonte protocol shows practical usefulness it is considered satisfactory in terms of reproducibility, assessment of evolution of pathology, and comparability between different settings, even though physician expertise is required [7,13]. Nevertheless, over the last few years, Niamey-protocol has been increasingly used in the quantification of PPF [7]

Recently, elastography methods, such as transient elastography (TE) and point shear wave elastography (pSWE) using acoustic radiation force imaging (ARFI), have been applied to assess liver fibrosis in some diseases, like chronic hepatitis C using US grading as gold standard [14−17]. These methods have also been considered promising for liver fibrosis assessment in patients with hepatitis B, non-alcoholic fatty liver disease, and post-transplant patients [15,18,19]. In addition, TE was recently evaluated to assess PPF induced by *S. mansoni* using US grading as gold standard [20,21].

As a friendly operator method, pSWE could be used as a complementary approach to assess PPF in schistosomotic patients. However, there is no systematic report of the literature on the use of pSWE for evaluation of PPF due to schistosomiasis mansoni. Therefore, we aimed to evaluate the performance of pSWE as a tool for predicting significant PPF in patients with schistosomiasis mansoni and to determine its best cutoff point, using US grading as gold standard.

## Materials and methods

### Patients

This cross-sectional study was conducted on adult patients with schistosomiasis mansoni, evaluated in the Gastroenterology outpatient clinic, Hospital das Clínicas of Universidade Federal de Pernambuco (HC-UFPE), from 2016 to 2017. Three hundred and ninety-six patients were eligible during the recruitment period. A total of 38 patients were excluded due to non-reliable pSWE results, characterized by an interquartile range of more than 30% of the median elasticity and a success rate of less than 60%. Among the 358 patients included, 222 (62.0%) were women, with a median age of 54 (ranging 18-92) years.

The diagnosis of schistosomiasis mansoni infection was based on clinical history of contact with fresh water sources from endemic areas, reports of prior treatment with praziquantel, along with US findings of PPF. Patients were classified into two groups according to the PPF patterns using Niamey’s classification: 86 patients with mild PPF (pattern C) and 272 patients with significant PPF (patterns D, E and F).

Patients with chronic viral hepatitis, fat liver disease, history of alcohol abuse, and other causes of liver disease, such as autoimmune hepatitis, drug-induced liver injury and haemochromatosis, were not included in this study.

### Abdominal ultrasound and liver stiffness measurement

All included patients were subjected to ultrasonography after to overnight fasting of about 8 hours, and were examined by the same experienced operator applying the Niamey classification, using a Siemens Acuson S2000 ultrasound system (Siemens Medical Solutions, Mountain View, CA, USA) with a 6C1 MHz transducer for both US and pSWE. Patterns of periportal fibrosis was registered according to the Niamey classification: pattern C (peripheral), D (central), E (advanced) and F (very advanced) [12,22].

After the US scan, the patients were submitted to pSWE, using the same machine. pSWE and measurements were acquired using Virtual Touch Quantification (VTQ) under standardized conditions on the right lobe (RL) of the liver, with the patient lying in a dorsal decubitus position, and the probe aligned along the intercostal space. The region of interest (ROI) was positioned into the hepatic parenchyma avoiding fibrosis tracts. The detection pulses measured the shear wave velocity, which was considered to directly relate to tissue stiffness. For each patient, valid shear wave speed (SWS) measurements were performed on the RL, and the results were expressed as the median value of the total measurements in meters per second (m/s). Reliable results were defined as those with an interquartile range (IQR) of less than 30% of the median value and a success rate of at least 60% [23].

### Ethics Statement

This study was approved by the Ethics Committee on Research Involving Human Subjects of the Health Sciences Center - Universidade Federal de Pernambuco (CCS-UFPE), Recife, Brazil (Approval no. 1.782.771/2016) and all patients were adults and signed terms of informed consent.

### Statistical analysis

The statistical analysis was carried out using the GraphPad Prism 6 software (GraphPad Software, Inc., La Jolla, CA) and MedCalc 15.2 software (MedCalc Software, Mariakerke, Belgium). A descriptive analysis including frequency, distribution, median and interquartile range was performed in both groups, and the D’Agostino & Pearson test was used to verify the normality of data distribution. A univariate analysis with χ^2^-test was used to compare categorical variables and Mann-Whitney test was used to compare two independent groups of continuous variables.

The performance of pSWE was assessed with receiver operating characteristic (ROC) curves. Optimal cut-off values for predicting significant PPF with best sensitivity and specificity were determined and positive (PPV) and negative predictive values (NPV) were calculated for these cut-off values. The area under the curve (AUC) was used to represent the accuracy of the predictions. A *p* value less than 0.05 was considered to be statistically significant.

## Results

The main characteristics of the patients included in the study are summarized in Table 1.

**Table 1.** Demographic, clinical and elastography parameters of 358 schistosomotic patients from Pernambuco/Brazil, according to periportal fibrosis pattern determined by ultrasonography, using Niamey classification.

|  | Periportal fibrosis pattern |  | *P* |
| --- | --- | --- | --- |
|  | Significant (D+E+F)<br>n=272 | Mild (C)<br>n=86 |  |
| Gender (n, %) |  |  |  |
| Male | 98(36.0) | 38(44.2) | 0.218<sup>a</sup> |
| Female | 174(64.0) | 48(55.8) |  |
| Age (years) * | 56(45-65) | 43.5(33.8-56.2) | **<0.001<sup>b</sup>** |
| Clinical form (n, %) |  |  |  |
| HSS | 220(80.9) | 5(5.8) | **<0.001<sup>a</sup>** |
| HS | 38(14.0) | 5(5.8) |  |
| HIS | 14(5.1) | 76(88.4) |  |
| SWS RL (m/s) * | 1.40(1.17-1.88) | 1.14(0.97-1.32) | **<0.001<sup>b</sup>** |

\*Median (interquartile range); <sup>a</sup> $\chi^2$ -test; <sup>b</sup>Mann-Whitney test;
HSS: hepatosplenic schistosomiasis; HS: hepatic schistosomiasis; HIS: hepatointestinal schistosomiasis; SWS RL: shear wave speed of the right lobe of the liver;

Measurements of SWS liver stiffness on the RL ranged from 0.66 to 4.27 m/s. The median values of liver stiffness assessed by pSWE, according to the PPF patterns, were: mild/pattern C of 1.14 m/s (IQR: 0.97-1.32); moderate/pattern D of 1.33 m/s (IQR: 1.03-1.69); advanced/pattern E of 1.40 m/s (IQR: 1.19-1.90); very advanced/pattern F of 1.53 m/s (IQR: 1.29-1.88). Some of the median liver stiffness values for PPF patterns showed substantial differences when compared: C vs D (p = 0.01), C vs E (p < 0.001), C vs F (p < 0.001), and D vs F (p = 0.014).

Among significant PPF patients, a higher medium age was identified (56 years, IQR: 45-65) and 64% were women. Comparing the clinical forms of schistosomiasis mansoni, higher frequencies of severe forms (hepatosplenic) were observed in significant PPF patients when compared with those of mild PPF, that showed higher frequencies of hepatointestinal forms, as shown in Table 1.

Results of liver stiffness measurement, in median values, by pSWE elastography revealed a substantial difference between patients with mild (1.14 m/s; IQR: 0.97-1.32) and significant PPF (1.40 m/s; IQR: 1.17-1.88), with a *p* value < 0.001, as shown in Table 1.

The best cutoff value of liver stiffness measurement for excluding significant PPF was equal or less than 1.11 m/s, with AUC of 0.719 and a sensitivity, NPV, and an overall accuracy of 80.5%, 40.5% and 71.2% respectively. Whereas, for confirming significant PPF the best cutoff value was above 1.39 m/s, with AUC of 0.719 and a specificity, PPV, and an overall accuracy of 86.1%, 92.0% and 59.2% respectively, as shown in Table 2 and Fig 1.

**Table 2.** Point shear wave elastography performance in the prediction of significant periportal fibrosis pattern in 358 schistosomotic patients from Pernambuco/Brazil, according to periportal fibrosis pattern determined by ultrasonography, using Niamey classification.

| SWS* | Periportal fibrosis pattern |  |  | Sensitivity |  | Specificity |  | PPV |  | NPV |  | A (%) |
| --- | --- | --- | --- | --- | --- | --- | --- | --- | --- | --- | --- | --- |
| | Significant (D+E+F)<br>n=272 | Mild (C)<br>n=86 | $p^{**}$ | (%) | 95% CI | (%) | 95% CI | (%) | 95% CI | (%) | 95% CI | |
| >1.11 | 219 | 50 | <0.001 | **80.5** | 75,3-85,0 | 41.9 | 31,3-53,0 | 81.4 | 76,2-85,9 | 81.4 | 30,2-51,4 | 71.2 |
| ≤1.11 | 53 | 36 |  |  |  |  |  |  |  |  |  |  |
| >1.39 | 138 | 12 | <0.001 | 50.7 | 44,6-56,8 | **86.1** | 79,9-92,6 | **92.0** | 86,4-95,8 | **92.0** | 29,1-42,5 | 59.2 |
| ≤1.39 | 134 | 74 |  |  |  |  |  |  |  |  |  |  |

\*Median values; \*\* $\chi^2$ -test;
SWS: shear wave speed; Sens: sensitivity; Spec: specificity; PPV: positive predictive value; NPV: negative predictive value; A: accuracy.

**Fig 1.**
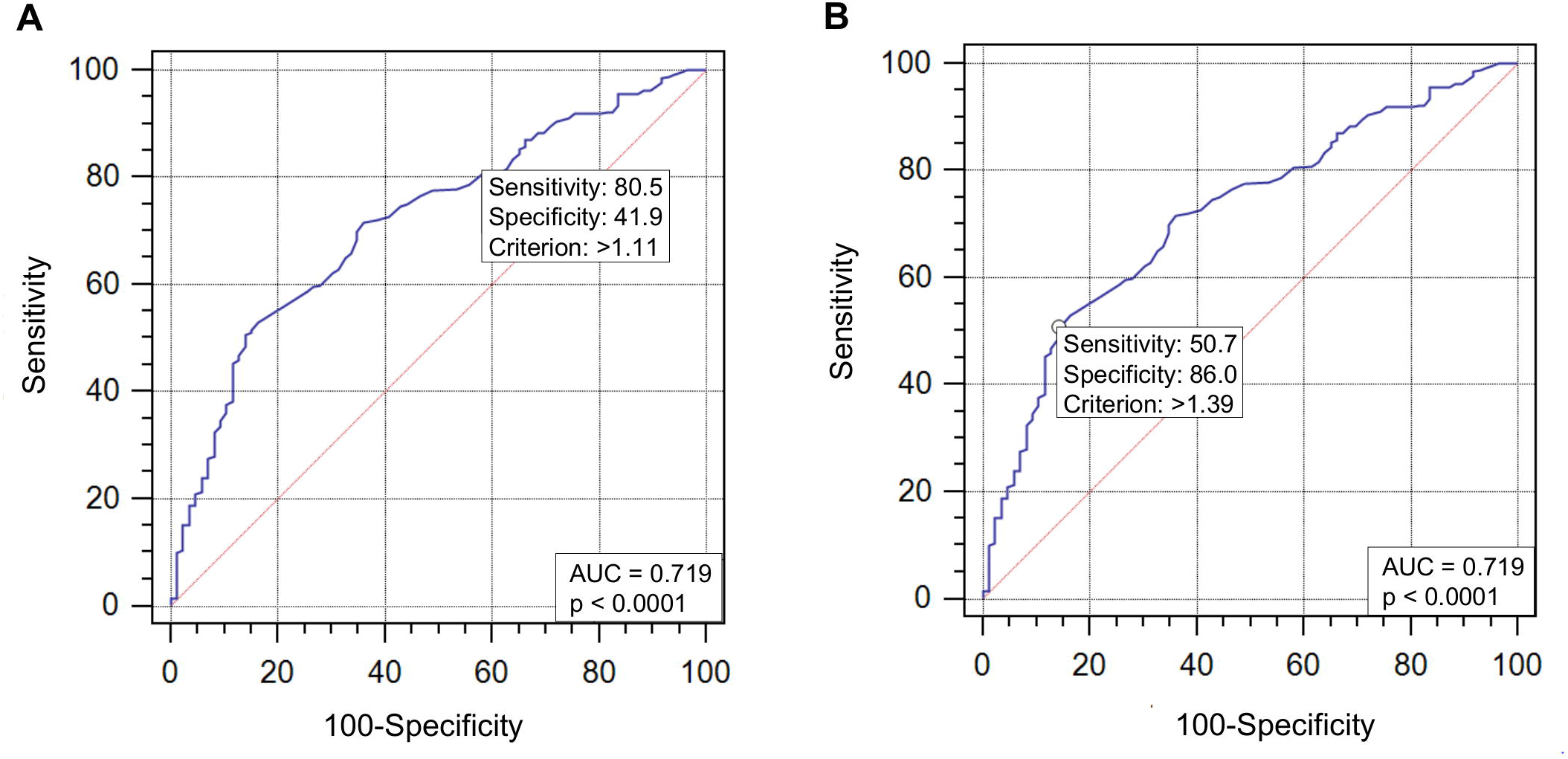
Receiver Operating Characteristic curve for shear wave speed of the right lobe of the liver to predict significant periportal fibrosis. (A) Cutoff point >1.11 with area under curve=0.719. (B) Cutoff point >1.39 with area under curve=0.719. Legend: AUC: area under curve.

Applying the cut-off values, 24.9% patients showed SWS values below or equal the lower cut-off point (≤ 1.11, n = 89) and among these patients, 40.5% were correctly classified, using US as gold standard. For 150 patients with SWS values above the upper cut-off point (> 1.39), 92.0% were correctly classified. In addition, 119 patients (33.2%) had indeterminate test values (between the lower and upper cutoff point: > 1.11 and ≤ 1.39) and among these, 68.1% had significant PPF and 31.9% had mild PPF.

## Discussion

According to liver stiffness measurement, our study showed that pSWE was a useful technique for confirming significant PPF. In addition, there were differences between the median liver stiffness values of mild pattern when compared to moderate, advanced and very advanced patterns, and also between moderate and very advanced patterns. Furthermore, the liver stiffness measurement was higher (p < 0.001) in the group with significant fibrosis-when compared with the mild fibrosis group.

Our results revealed that the best cutoff value for excluding and confirming significant PPF was equal or less than 1.11 m/s and above 1.39 m/s, with a AUC of 0.719 with sensitivity of 80.5% for the lower cutoff and specificity of 86.1% for the higher cutoff. Accordingly, the method showed better performance to confirm significant PPF, since more than 90% of patients were correctly classified.

Similarly, Veiga et al., in a multicentric study that included patients from Rio de Janeiro, Brazil, compared liver stiffness among hepatosplenic schistosomiasis, hepatitis C virus-cirrhotic, and control patients using TE. They concluded that liver stiffness evaluation may be a useful tool to differentiate portal hypertension related to cirrhosis from that of hepatosplenic schistosomiasis [21]. In this study, however, no association was detected between Niamey PPF patterns and TE evaluation. The lack of association between these parameters in this study might have been caused by the small sample size, predominant inclusion of patients with mild PPF patterns (patterns B and C) and lack of patients with more advanced patterns (patterns E and F).

On the other hand, Shiha et al. reported that TE would not be useful to diagnose liver fibrosis and esophageal varices in patients with pure schistosomiasis [20]. Nevertheless, these authors, did not applied the Niamey classification, and their paper aimed to compare patients with and without esophageal varices and splenomegaly with TE results.

Moreover, since schistosomiasis PPF is not diffuse by the liver parenchyma, TE evaluation does not allow ROI to be placed in a suitable area, which is between the periportal fibrosis septa. On the other hand, pSWE enable a completely view of the liver parenchyma and ROI positioning to improve liver stiffness evaluation in this area.

US scan has become the most consolidated tool for evaluation of PPF in endemic areas because it shows practical usefulness, satisfactory interobserver agreement, and enables data comparison on international scale [6−8,21]. Despite the widespread use of US for diagnosing and monitoring changes caused by PPF, its use has some limitations, since it is a subjective method and requires physician expertise to apply the WHO Niamey-Belo Horizonte protocol.

In our region, Northeast Brazil, which is endemic for schistosomiasis mansoni, the US scan is widely used in clinical practice, mainly in the rural zone and for scientific research in the field. Consequently, with the recent advent and widespread use of liver elastography, we aimed to evaluate the performance of pSWE as complementary tool for assessment and staging of PPF in schistosomotic patients. This method was chosen since it is more easily applied than Niamey US protocol, and, could enhance the diagnosis of those patients with advanced PPF, who need greater care and supervision. Moreover, it would provide a foundation for further studies to assess the presence and severity of portal hypertension and prediction of esophageal varices using pSWE and TE [21,24−26].

pSWE is a promising US-based method and it has been currently used to assess the severity of liver fibrosis in patients with chronic viral hepatitis, showing accuracy similar to TE [15,16]. In addition, pSWE has also showed promising results in patients with non-alcoholic fatty liver disease and non-alcoholic steatohepatitis [27]. As advantages pSWE is a technique that can be integrated into routine ultrasound protocols, it can be used in patients with ascites, require less sophisticated skills from the operator, and it is also possible to determine optimal region of interest (ROI) placement, avoiding structures such as focal lesions, large blood vessels and heterogeneous areas [15,28].

Nevertheless, there are only few studies evaluating elastography in schistosomotic patients, and schistosomiasis mansoni monoinfection has not been evaluated by pSWE so far. Accordingly, in studies evaluating coinfected patients with PPF and chronic hepatitis B [29] or C [25,26,30], the liver stiffness measurement using pSWE could be overestimated. Moreover, patients with schistosomiasis mansoni are usually excluded from elastography analyzes to assess the severity of liver fibrosis in patients with viral hepatitis [31,32].

Overall, this study showed some limitations considering that US and pSWE were performed by the same operator. Nonetheless, the pSWE assessment was performed after the US scan, avoiding or reducing operator subjectivity interference. Additionally, further studies comparing US and pSWE results of different operators and analyzing inter and intraobsever variability, would provide better evidence about pSWE reproducibility for predicting PPF.

In the future, new portable US devices capable of performing pSWE can be developed, allowing this technique to be applied to schistosomotic patients in endemic areas, especially in the field. Therefore, it will be possible to improve these patients’ diagnosis and prognosis without the need of a trained examiner.

In conclusion, pSWE elastography was able to differentiate significant from mild PPF, with better performance to predict significant PPF. Therefore, pSWE is a potential tool for noninvasive assessment of disease severity in patients with schistosomiasis mansoni.

## Acknowledgements

The authors thank Jair Matos from Siemens Healthineers for technical assistance.

## Author Contributions will

○ *Conceptualization: ADB AD EPAL ALCD*
○ *Data Curation: JCS CMMV VRSJ*
○ *Formal Analysis: JCS ADB EPAL ALCD*
○ *Funding Acquisition: AD HD SMLM ALCD*
○ *Investigation: ADB RSL ALCD*
○ *Methodology: JCS ADB EPAL ALCD*
○ *Writing – Original Draft Preparation: JCS*
○ *Writing – Review & Editing: ADB EPAL ALCD*

## SUPPORTING INFORMATION LEGENDS

S1 Checklist: STROBE Checklist

